# Rapid memory reactivation at movie event boundaries promotes episodic encoding

**DOI:** 10.1101/511782

**Authors:** Marta Silva, Christopher Baldassano, Lluís Fuentemilla

## Abstract

Segmentation of continuous experience into discrete events is driven by rapid fluctuations in encoding stability at context shifts (i.e., event boundaries), yet the mechanisms underlying the online formation of event memories are poorly understood. We investigated the neural spatiotemporal similarity patterns of the scalp electrophysiological (EEG) activity of 30 participants watching a 50 min movie and found that event boundaries triggered rapid reinstatement of the just-encoded movie event EEG patterns. We also found that the onset of memory reinstatement at boundary onset (around 1500ms) was preceded by an N400-like ERP component, which likely reflects the detection of a context switch between the current and just-encoded event. A data-driven approach based on Hidden Markov Modeling allowed us to detect event boundaries as shifts between stable patterns of brain EEG activity during encoding and identify their reactivation during a free recall task. These results provide the first neurophysiological underpinnings for how the memory system segments a continuous stream of experience into episodic events.

## Introduction

Memory systems transform the stream of our continuous experience into a sequence of episodic memory units to be recalled in the future. While extensive research has been conducted to understand how the brain supports the formation of discrete, brief novel information, it is only recently that psychologists and neuroscientists have started exploring the mechanisms that account for episodic memory formation during a continuous stream of experience.

A widely accepted view is that we naturally segment continuous experience into events, and that event boundaries are driven by moments in time when prediction of the immediate future fails (Zack et al. 2007) or by fluctuations in contextual stability (Clewett & Davachi 2017). Segmentation affects not only our perception of the experience, but its subsequent organization in long-term memory (Kurby and Zacks, 2008; Radvansky, 2012; Sargent et al., 2013), such that elements within an event are bound together more cohesively than elements across events (Ezzyat and Davachi, 2011; DuBrow and Davachi, 2013 and 2014; Horner et al., 2016). Human neuroimaging studies using naturalistic video clips have set important findings that align well with these behavioral findings. They have shown that a distributed network of brain regions comprising the hippocampus and neocortex are involved during event segmentation and that their dynamics during encoding provide a basis for how we parse the temporally evolving environment into meaningful units. They revealed that the brain organizes the ongoing input into episodic events by detecting changes in the stability of activity patterns. Stable patterns of activity at higher-level brain regions during encoding are thought to maintain a stable event representation in spite of fluctuations in the ongoing sensory input (Chen et al., 2017; Baldassano et al., 2017). Shifts in stability that coincide with perceived boundaries induce a neural response at the hippocampus (Ben-Yakov et al., 2011, 2013 and 2018; see Bulkin et al., 2018 for similar findings in rodents) and the degree to which hippocampal activity at boundaries couples with cortical patterns of activity predicts pattern reinstatement during later free recall (Baldassano et al., 2017), thereby indicating that the hippocampus may be responsible for binding cortical representations into a memory trace online during encoding (McClelland et al., 1995; Moscovitch et al., 2005; Norman and O’Reilly, 2003). However, an important question remains unanswered: which neural mechanisms support the binding of the encoded information of an event upon boundary detection? And more importantly, how can we investigate these neural mechanisms in ecologically valid circumstances that can inform us about their nature in real life environments?

To address this issue, we recorded scalp brain electroencephalography (EEG) while 30 participants watched a single 50-min movie clip and asked whether time-resolved fluctuations in neural similarity elicited during movie viewing reflected event segmentation. Leveraging the fine-grained temporal resolution of the EEG signal, we then tested the hypothesis that moments in time after perceived event boundaries during movie viewing would exhibit reactivation of the just-encoded episode, and that this reactivation would promote consolidation of the encoded event into long-term memory. Indeed, the reactivation of encoded episodes upon boundary detection would be in line with animal research using EEG recordings showing that memory replay of the just-encoded event promoted its memory formation and consolidation (Carr et al., 2011) and with recent EEG research in humans that showed that memory reactivation at picture boundaries during sequence encoding promoted a linked memory representation across events (Sols et al., 2018). The extent to which boundary-triggered memory reactivation impacted memory formation during movie viewing would offer valuable insights into how the brain shapes the unfolding experience into memory under ecologically-valid situations.

## Results

### Event segmentation and perceived event boundaries

Six external participants, who did not take part in the electrophysiological recording session of the study, were asked to watch the first episode of the Sherlock TV series (Baldassano et al., 2017; Chen et al., 2017), which lasted 50 min. Using the standard event segmentation approach from Newtson, 1973 and Zacks et al., 2010 (also used in Baldassano et al., 2017, Chen et al., 2017; and Ben-Yakov and Henson, 2018), participants were requested to annotate with precision the temporal point at which they felt ‘‘a new scene is starting; these are points in the movie when there is a major change in topic, location or time.” Participants were also informed that each event should be between 10 seconds and 3 minutes long and we asked them to write down a short title for the event.

Temporal points at which at least three external raters coincided in annotating a boundary were taken as indicative of a “real” event boundary in the movie (see Methods). This approach resulted in an event segmentation model of 38 event episodes (Figure 1a), which was consistent with the range found in our previous study (Baldassano et al., 2017).

**Figure 1.**
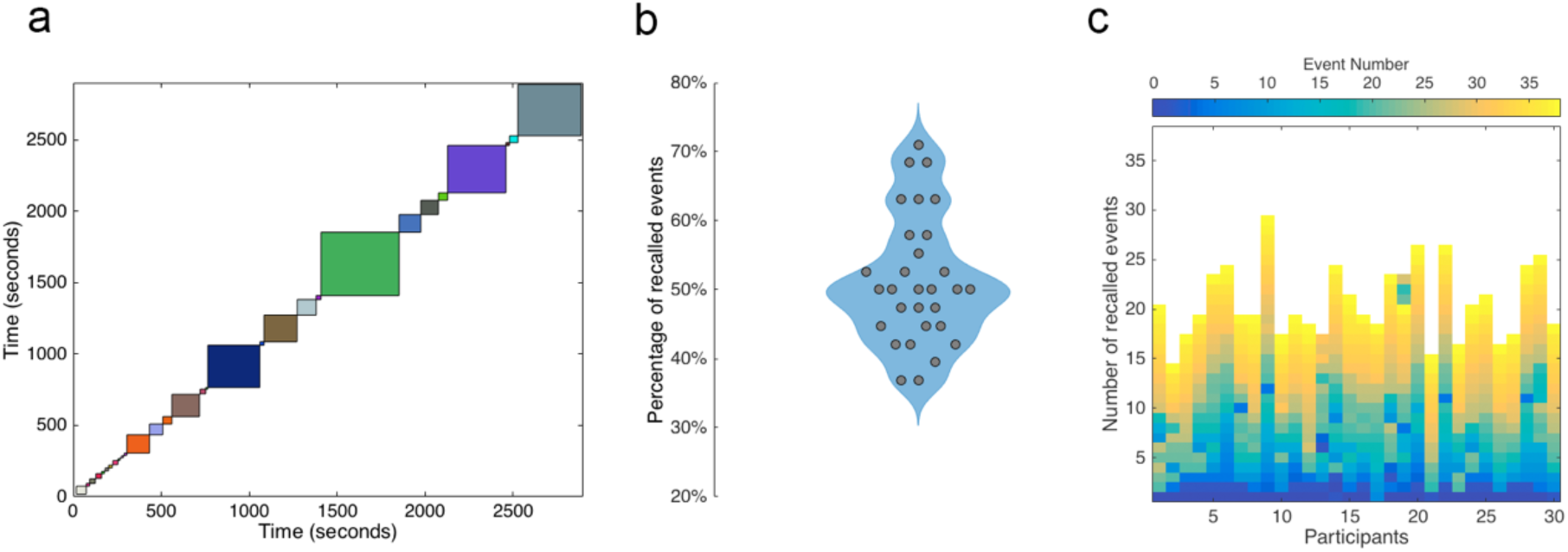
Event Segmentation Model and memory performance. (**a**) Schematic representation of the event segmentation model derived from human annotations. Each color-coded square denotes events during the 50-min movie and start/end of each event represents the boundary time points. (**b**) Proportion of events that were later recalled by the participants in our sample (N = 30). (**c**) Color-coded temporal order distribution of movie events that were recalled in the free recall task for each participant.

### Movie free recall

We then recruited 30 different healthy participants to participate in an EEG recording session, during which they were asked to watch the same movie. After 15 min of rest, all subjects in this dataset were asked to retell the story they had just watched, without any cues or stimulus. EEG was also collected during this time, and verbal recall was recorded through an audio recorder for later analysis.

Participants’ memory accuracy indicated they were able to recall 51.4% of the encoded events on average (STD = 9.2%) (Figure 1b). Importantly, we also found that the temporal order of the episodic events at encoding was preserved at recall (Figure 1c; Supplementary Figure 1), thereby replicating previous results (Chen et al. 2017) that free recall tends to preserve the temporal structure of the encoded memories.

### Event segmentation model on the EEG data

Next, we tested whether patterns of EEG activity elicited by the 50-min movie exhibited the event structure hypothesized by our model (periods with stable event patterns punctuated by shifts between events). To address this issue at individual level, we computed a point-to-point spatiotemporal similarity analysis of the EEG data during the 50-min movie-watching and calculated the degree of similarity values within each of the events (defined by the human-annotated event boundaries) (Figure 2a). To statistically assess the extent to which the EEG data fit the model, we averaged the spatiotemporal similarity values within each of the 38 events and tested this value against a null distribution generated by running the same analysis 1000 times with a shuffled temporal order distribution of the events (Figure 2b; see Methods). This analysis revealed that 22 out of the 30 participants in our sample showed a higher degree of similarity values within events from the real segmentation model as compared to their individual correlation value cut-off (alpha = 0.05) by the null distribution (Figure 2c) and that this was significant at group level (p < 0.05; Figure 2d).

**Figure 2.**
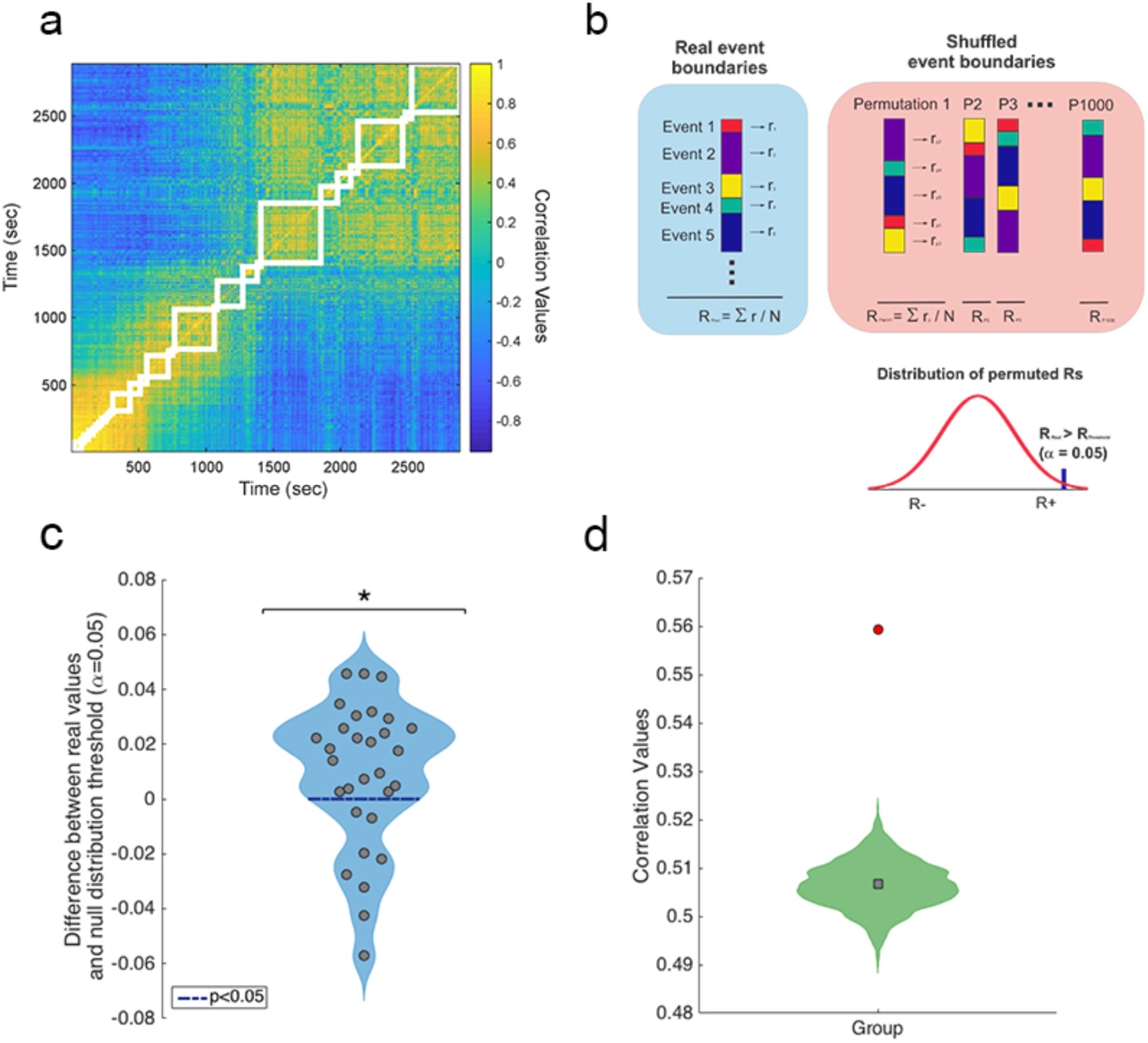
EEG neural patterns during movie watching and event segmentation model. (**a**) A temporal correlation matrix was generated from raw EEG data for each of the participants (an example of one selected participant is depicted in this figure). The event segmentation model from human-labeled boundaries is overlaid in white. (**b**) For each participant, the event segmentation model was used to calculate the averaged correlation values for pairs of timepoints within each event. A null distribution of correlations was obtained for random event boundaries by shuffling the order of the events of the segmentation model 1000 times. (**c**) Single-participant distribution of the difference between the real within-event correlations and α<0.05 thresholds from the null distribution. *denote the results were significant at group level (p < 0.05). (**d**) The red circle shows the true participant average; green histogram shows the null distribution of the participant average; grey square shows mean of the null distribution.

### Shared event neural patterns across participants during movie-watching

Having shown that EEG patterns of neural activity were structured according to a general event segmentation model during movie-watching, we then tested the prediction that within-event EEG patterns should be shared across individuals (Chen et al., 2017). To address this question empirically, we computed Movie-Movie correlations by comparing patterns of each event from one participant with the movie pattern for the same event averaged across the remaining participants. An averaged correlation value was obtained for each participant and its statistical significance was assessed by comparing it to a random distribution obtained by shuffling the event order on the left-out participant. Confirming previous findings on fMRI data (Chen et al., 2017), we found that almost all of the participants (29 out of the 30) showed high degree of shared similarity EEG patterns with the group sample (p < 0.05 at group level) (Figure 3a and b).

**Figure 3.**
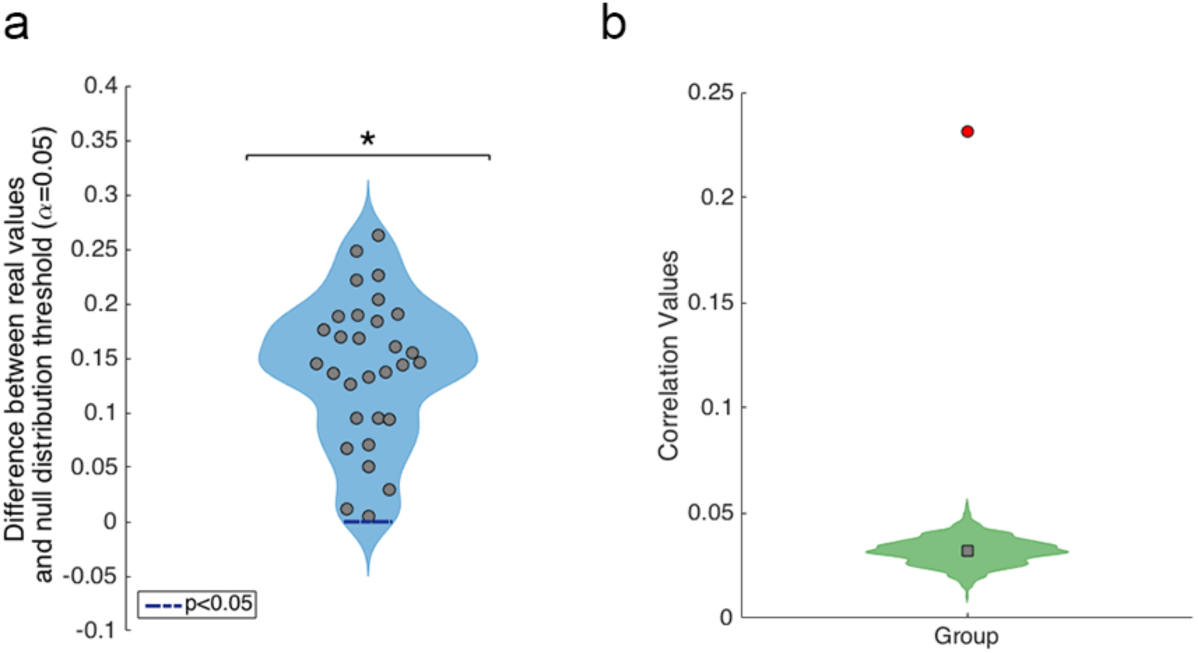
Between-participant pattern similarity during movie viewing. (**a**) Inter-subject correlation value derived from correlating the patterns for each event in each individual with the corresponding event patterns in the rest of the group, compared to an α < 0.05 threshold from the null distribution. *denote the results were significant at group level (p < 0.05). (**b**) The red circle shows the true participant average; green histogram shows the null distribution of the participant average; grey square shows mean of the null distribution.

### EEG pattern similarity within and across events separated by boundaries

An important assumption derived from event segmentation theory is that patterns of neural activity elicited within an event should be more stable than neural patterns across events, thereby indicating that event neural representations change when boundaries are detected. To test this prediction in our data, we ran a point-to-point spatial similarity analysis throughout EEG segments of -10 sec to 10 seconds of averaged EEG trials around the boundary time point. The long EEG segments were then split into EEG epochs of 5 seconds each, thereby allowing us to examine the extent to which similarity values were higher for neural responses within events. More concretely, the similarity analysis was performed between three different pairs of temporal intervals in the data: pre-boundary time intervals (−10 to -5 sec and -5 to 0 sec to the boundary), between-event time intervals (−5 to 0 and 0 to 5 sec to the boundary) and post-boundary time intervals (0 to 5 sec and 5 to 10 sec to the boundary) (Figure 4a). The resulting similarity values for each condition and subject were then averaged and differences were tested by means of a repeated measures ANOVA. Notably, the results from this analysis revealed that similarity values differed between conditions (F(2,58)=13.94, p < 0.001). Post-hoc paired t-test showed that within event similarity, both pre-boundary and post-boundary, were higher than between event similarity (t(29) = 4.44, p < 0.01 and t(29) = 3.46, p < 0.01, respectively) and that similarity values within pre and post-boundary conditions were statistically equivalent (t(29) = 0.83, p = 0.41) (Figure 4b).

**Figure 4.**
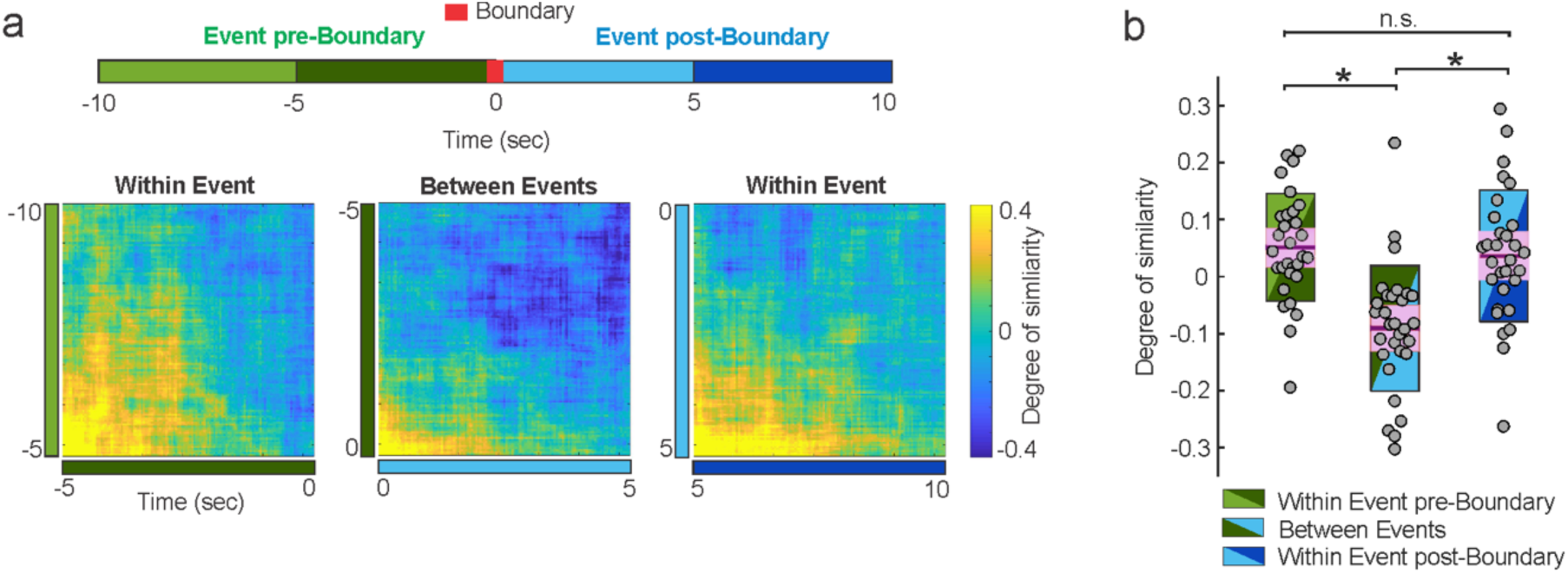
Neural pattern similarity within and across events during movie-watching. (**a**) A time-resolved similarity analysis was calculated for pairs of samples over 20 seconds around boundaries, grouped based on whether the two samples fell before the boundary, on both sides of the boundary, or after the boundary. (**a**) Time-resolved degree of similarity averaged over participants for EEG activity within events before the boundary, across events separated by boundaries and within events post-boundaries. (**b**) Participant’s degree of similarity for each of the event conditions depicted in (b). For all boxplots, the central mark is the median, the edges of the box are the 25^th^ and 75^th^ percentiles. * denotes p < 0.05.

### Boundaries trigger rapid reactivation of the just encoded event during movie viewing

Leveraged by previous findings indicating that explicit context shifts triggered rapid reinstatement of the just-encoded picture event sequence and that such reinstatement at boundaries promoted the formation of long-term memory for that event (Sols et al., 2017), we tested the prediction that neural reactivation may also support memory formation of the just-encoded event during much more subtle transitions between events during movie-watching, providing converging evidence that memory reinstatement at event boundaries facilitates the storage of events into long-term memory. To address this hypothesis, we computed a neural similarity analysis between EEG data epochs of 10 seconds preceding and following boundary timepoints and compared the resulting similarity values for events that were later recalled in the free recall task with events that were later forgotten. This analysis revealed that patterns ∼1.5 seconds post-boundary were significantly more similar to patterns ∼5-10 seconds pre-boundary when these pre-boundary events were later recalled (Figure 5a). These findings provide evidence, for the first time, that neural reactivation is a mechanism supporting the formation of event episodic memories upon boundary detection during a continuous stream of stimuli.

**Figure 5.**
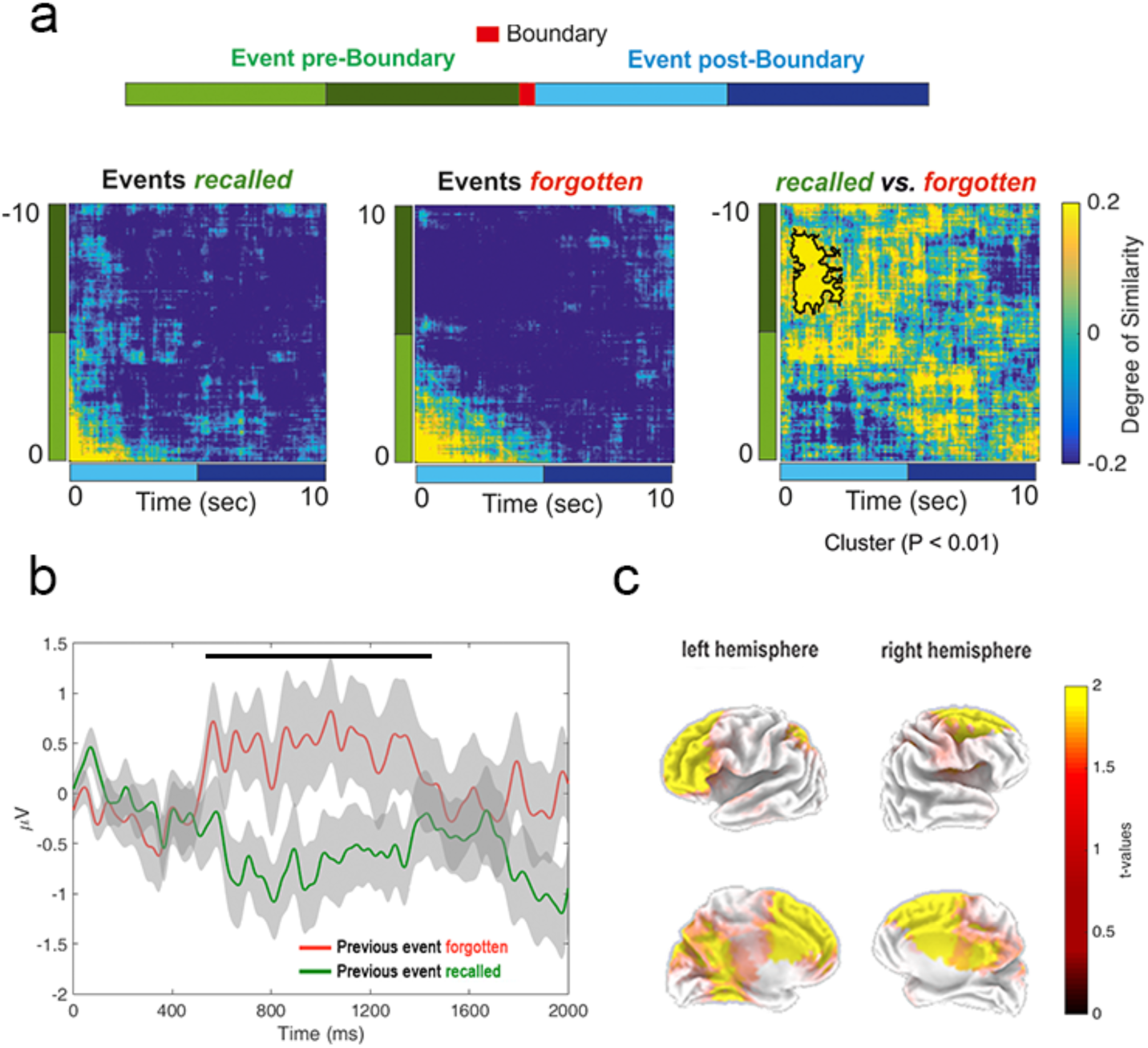
Rapid neural reinstatement and evoked response at boundaries during movie-watching. (**a**) Time-resolved degree of similarity across event boundaries that followed events that were later recalled or forgotten. Difference between similarity values for the two conditions is depicted on the right. Statistically significant (p < 0.05 at cluster level) greater similarity was found across events for EEG at around 1.5 seconds at boundary onset (indicated with a black thick line). (**b**) Event Related Potentials (ERPs) elicited at boundary onset during movie watching as a function of whether the previous event was recalled or forgotten in the subsequent recall task. Thick lines depicted the averaged ERPs over the 29 scalp electrodes across participants and the shaded area represents standard error of the mean of the participants’ sample. The thick black line depicts the timing of the significant cluster between ERP conditions. (**c**) Brain sources of the ERP difference observed at boundary onset between recall and forgotten conditions.

### Neural responses preceding neural reactivation at boundaries

Though memory reactivation was found to take place rapidly upon boundary onset (i.e., ∼1.5 seconds) in our study, research on Event-Related Potentials (ERPs) have also revealed the existence of even faster event-locked neural responses predictive of successful encoding (i.e., within the first second of event onset) (Friedman and Johnson, 2000). One such an example in the context of current study is the N400, which is a well-known negative ERP component appearing around ∼600 ms after stimulus input that assesses semantic memory states, with the amount of N400 amplitude variation revealing how much of the information elicited by that stimulus is congruent with the already active one (Kutas and Federmeier, 2011). Thus, given that memory formation of a meaningful event may depend on the ability to perceive a contextual shift at a boundary (i.e., segmentation; Kurby and Zacks, 2008), we asked whether perceived boundaries following events that were later recalled triggered an N400-like component within a time window preceding the onset of neural reactivation. To address this issue, we compared the ERPs time-locked to boundaries following events that were later recalled and forgotten and we found that these two conditions showed a differential ERP in a time-window of 600-1400ms after boundary onset, being more negative in polarity after boundaries following events that were later recalled (Figure 5b). Importantly, this ERP difference was not observed when the same analysis was performed at neural responses time-locked to boundaries *preceding* later recalled and forgotten events (Supplementary Figure 2) and these findings could not be attributed to a distinct proportion of events that were later recalled or forgotten following the boundary event either (Supplementary Table 1), indicating that the memory formation processes associated to memory formation had a retrospective nature when boundaries were detected. Furthermore, a source brain analysis revealed that activity from frontal, parietal and medial temporal regions were involved in the ERP differences between conditions (Figure 5c), matching brain regions found to be associated with event boundary segmentation in our previous fMRI study (Baldassano et al., 2017). Altogether, these findings suggest that event boundaries trigger a sequence of neural mechanisms associated with first signaling a shift between previous and current ongoing event information followed by a rapid reactivation of the just encoded event.

### Neural reactivation during free recall

An intriguing finding in our previous fMRI study using the same movie was that the elicited patterns of neural activity associated with event segmentation during encoding were later reinstated during free recall (Baldassano et al., 2017). The extent to which these findings could be replicated using electrophysiological recordings may be relevant to open new venues for examining the neural mechanisms supporting event structure reinstatement patterns. To address this possibility, we adapted the approach implemented in our previous study based on Hidden Markov Modelling (HMM) to the present EEG study (see Methods). Briefly, the HMM approach implements a data-driven segmentation search and returns the most probable division of a given signal to a given number of events. An important advantage of the HMM in the context of this study is that it provides a data-driven solution for how the ongoing pattern of neural activity may be sequenced into a given number of events. This is an attractive approach as it allows searching for the existence of patterns of neural activity related to complex event sequence structure in a flexible manner, as the algorithm can be applied to brain signals of different length. This is particularly relevant in the context of a free recall task, as in the current study, given that total recall length and per-event recall time length varied across participants and within participants respectively (Figure 6a). Thus, for each participant, the HMM was used to estimate a 38-event segmentation of the continuous EEG data acquired during recall that most closely corresponded to the 38 neural event patterns elicited during movie-watching. If, according to our previous findings using fMRI (Baldassano et al., 2017), participants’ recall involved the reinstatement of neural patterns during encoding, we would expect event-elicited EEG activity during encoding and HMM-derived event-elicited EEG activity at recall to be very similar. To measure this for each individual, we correlated the EEG patterns elicited during encoding and recall and tested whether the resulting correlation value was higher than in a null distribution obtained by shuffling the order of events between encoding and recall. We found that 27 participants, out of the 30 in the sample, showed that real movie-recall correlation was higher than the threshold (p < 0.05) and that this proportion of significant findings in our sample could not be attributed to chance at group level (p < 0.05 at group level) (Figure 6b and c). This result extends previous fMRI findings (Baldassano et al., 2017), demonstrating that memory recall is supported by the reinstatement of the electrophysiological patterns elicited during movie-watching.

**Figure 6.**
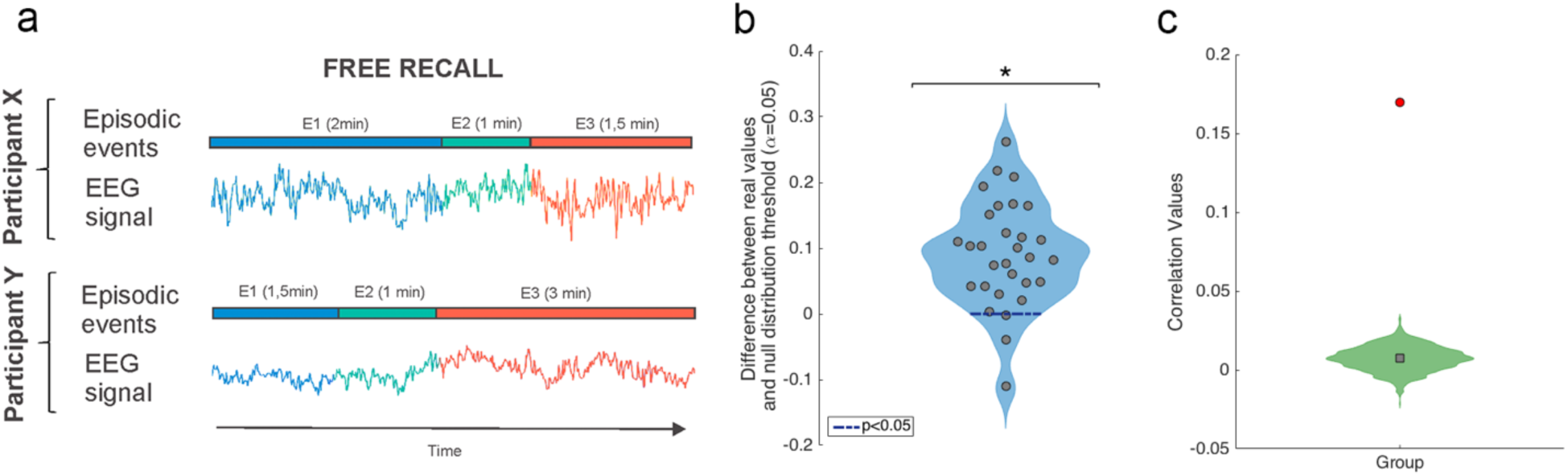
Neural reinstatement at recall. (**a**) Illustration of how two participants’ recall lengths varied for the same three events. Using an HMM approach, we searched for reinstatement of EEG neural event patterns in spite of these differences’ length. (**b**) EEG correlations between event segmentation model patterns during movie watching and recall activity derived from HMM-estimated events were statistically significant for 23 out of 30 participants of the sample. The difference between real correlation and α < 0.05 threshold set by randomly permuting the order of the events at encoding is displayed for each individual. * indicates the results were significant at group level (p < 0.05). (**c**) The grey circle shows the true participant average; green histogram shows the null distribution of the participant average; red square shows mean of the null distribution.

## Discussion

Our results provide the first evidence of electrophysiological signatures related to how event segmentation during movie viewing shapes memory formation. They show that patterns of neural activity recorded from the scalp EEG while viewing a 50-min movie fit with an event segmentation model of episodic events punctuated by rapid transitions of content (i.e., boundaries). We observed that these event-specific patterns of neural activity were reinstated at later recall, thereby corroborating the idea that the event segmentation process shaped memory formation of a continuous stream of stimuli into a structured memory representation that can be accessed long-term. Leveraged by the fine-grained temporal resolution of the EEG data, we showed that event memory formation during movie viewing was mediated by the extent to which it was rapidly reactivated at event boundaries and that memory reactivation was preceded by an N400-like ERP component time-locked to the boundary, which likely reflect the effective detection of a context switch between the current and just-encoded event. These findings indicate that the successful encoding of an episode is regulated by two neural mechanisms that act sequentially within the first ∼2 seconds after an event boundary.

Why would memory reinstatement be advantageous during the encoding of a continuous stream of stimuli? Though event segmentation provides a framework to examine how continuous experience can be chunked into a set of discrete episodes in memory through the detection of event boundaries, it does not account for how this sequence of episodes can be integrated into a memory structure that preserves the temporal structure during later recall. Memory reinstatement at event boundaries may represent a way to promote temporal event memory organization across boundaries as it may serve to promote the strengthening, or chunking, of that just-encoded event but it also may help promote binding across boundary episodes as a result of the contemporaneously co-activation of the past and present events (Sols et al., 2017). The extent to which memory reactivation at event boundaries serves to promote the encoding of unique events into memory, the integration of different events into a temporally organized memory structure or both is difficult to disambiguate in our study as participants memory accuracy was obtained through a free recall task, which relies on retrieval processes heavily dependent on clustering properties of the encoded material, such as semantic similarity or temporal proximity (Polyn et al., 2009).

Speculatively, it could be argued that memory reactivation at event boundaries could theoretically represent a way to account for how different event episodes that shared contextual semantic properties can be integrated. In support of this hypothesis, previous fMRI studies have shown that temporally extended events sharing contextual information were evaluated as if they appeared closer in time during recall and that this was related to increased hippocampal similarity between these events during encoding (Ezzyat and Davachi, 2014). Interestingly, this effect was only observed for when events that shared contexts were separated by event boundaries during their encoding, suggesting the possibility that neural mechanisms triggered at boundaries were at least partially responsible for memory integration (e.g., memory reactivation). Another set of research studies have emphasized the relevance of memory reactivation to explain how different episodes are integrated as a function of the degree of their overlapping content to allow generalization (Schlichting and Preston, 2015). These studies have shown that memory reactivation is elicited when elements of the experience partially mismatch with stored memory representations, supporting integrative encoding online (Shohamy and Wagner, 2008).

An open question is which specific mechanisms trigger memory reinstatement at event boundaries. Advancing on this issue is not trivial given the diverse ranges of stimulus used in event cognition literature (e.g., text narratives (Zwaan, 1996), short video clips (Ben-Yakov et al., 2011 and 2013) or item sequences (Dubrow and Davachi, 2013)). In an attempt to accommodate the literature on this topic, Clewet and Davachi (2018) suggested that event boundaries represent moment-to-moment fluctuations in external and internal contextual states during continuous encoding. In our study, such fluctuations could be understood as moments in time when an internal representation derived from an accumulated contextual encoding suddenly shifts at the start of a novel scene with a change in spatial location, characters or goals. Interestingly, we found that these moments in time were followed by an N400-like ERP component, specifically for when the previous event was later recalled but not for those that were forgotten. The fact that this specific ERP response preceded the onset of the memory reactivation after an event boundary lends support to the notion that it could reflect a neural process that may be necessary to trigger neural reactivation. However, this idea should be taken cautiously as our data do not provide a quantifiable dependency between the two other than differences in their temporal onset. Nevertheless, our results offer valuable testable predictions for future research on how perception and memory interact online during the encoding of ecologically-valid stimulation protocols.

In a series of fMRI experiments, Ben-Yakov et al. (2011, 2013) revealed the importance of studying event offset brain activity in humans at the end of movie clips to understand how episodic memories are formed during the stream of a continuous audiovisual stimulation. These studies, together with those studying abrupt switches between stimulus category and task (DuBrow and Davachi, 2016), and recent studies of event boundaries in movies (Ben-Yakov et al., 2018; Baldassano et al., 2017), offered converging evidence for the specificity and sensitivity of the coupling of the hippocampus to event boundaries during movie viewing. In the current study, we found that the brain sources of the N400-like ERP associated with memory formation at boundaries of the just-encoded event episode included frontal, parietal and medial temporal lobe regions. These regions highly overlapped with brain regions found in our previous fMRI findings of hippocampally-linked event boundaries (Baldassano et al., 2017). The similarity of these sources represents strong evidence that our approach was suitable for identifying the engagement of the same brain network and informing about the temporal properties of their engagement using non-invasive electrophysiological recordings.

In naturalistic scenarios, the study of the recollection of memories of one’s past may vary substantially across individuals and within subjects as a function of task contexts. This creates an important challenge in our search for the neural underpinnings supporting the remembering of autobiographical memories. The implementation of data-driven modeling approaches, such as the ones offered by HMM used in the current study, may foster interesting possibilities in this endeavor. Indeed, being able to identify the reinstatement of memory events from a 50-min movie viewing using HMM extends previous fMRI findings (Baldassano et al., 2017). Our work is, however, the first to show that HMM can be used to model electrophysiological signals, thereby proving its usefulness to test predictions of how perception and memory are supported by the brain that rely on fine-grained temporal dynamic resolution, such as whether neural oscillations at different frequencies support the hierarchical structure found in Baldassano et al. (2017), or whether specific oscillatory activity related to memory formation and retrieval (i.e., theta frequency band, 3-8Hz) play a role at event boundaries.

Understanding how memories are formed and structured in real life requires the characterization of neural mechanisms that take place online, during the ongoing encoding of a continuous naturalistic stimuli, as our experience unfolds over time. Investigating how memories are formed during audiovisual narratives such as long movie clips may provide a valuable approach to bring testable predictions derived from animal and theoretical neuroscience into real life settings. The current experiment assessed whether memory reinstatement, a neural mechanism critical for memory formation and consolidation, took place under these ecologically-valid experimental circumstances. By showing that this is the case, our findings offer valuable insights into how the brain shapes the unfolding experience into long-term memory that can be generalized to real life.

## Methods

### Participant sample

Thirty-three Spanish speakers (30 right-handed, 20 females, age range 18-46, mean = 22) participated for pay (10€/h). Participants were recruited from the University of Barcelona and the broader community. All participants were healthy and did not consume psychoactive substances. Informed consent was obtained from participants in accordance with procedures approved by the Ethics Committee of the University of Barcelona. Data from two participants were discarded due to falling asleep during the experiment, and one due to too much muscular artifact in the data. Thus, the final sample of participants included in the study was thirty.

### Experimental Design

Our primary dataset consisted of 30 participants who watched the first 50 minutes of the first episode of BBC’s *Sherlock* in Spanish and were then asked to freely recall the episode without cues, while being recorded using an audio recorder. The audio files were later analyzed in order to access participants’ length and richness of the recall, with total recall times ranging from 6 min to 44 min (and a mean of 15 min). At the beginning of the movie, a 30 s audiovisual cartoon (“Let’s All Go to the Lobby”) was presented to set participants attention. The experimental design was implemented on ePrime 2.0 (Psychology Software Tools, Inc.)

### EEG recording and preprocessing

EEG was recorded using a 32-channel system at a sampling rate of 500 Hz, using a BrainAmp amplifier and tin electrodes mounted in an electrocap (Electro-Cap International) located at 29 standard positions (Fp1/2, Fz, F7/8, F3/4, Fc1/2 Fc5/6, Cz, C3/4, T3/4, Cp1/2, Cp5/6, Pz, P3/4, T5/6, PO1/2, O1/2) and at the left and right mastoids. An electrode placed at the lateral outer canthus of the right eye served as an online reference. EEG was re-referenced offline to the right and left mastoids. Vertical eye movements were monitored with an electrode at the infraorbital ridge of the right eye and an independent component analysis (ICA) was run on MATLAB’s EEGlab toolbox (Delorme et al., 2004) to correct for eye movements and remove extremely noisy components (no more than 6 components were removed).

Using the EEGLAB toolbox a low pass filter of 20Hz was applied in order to reduce the presence of muscular artifacts (Perez et al., 2017). Data corresponding to encoding (e.g. movie viewing) and recall was down-sampled by segmenting the EEG into bins of averaged data from one hundred sample points (i.e., 200 milliseconds). We choose this interval as a compromise between preserving data structure and reducing computational time in our analyses.

### Event boundary annotations by human observers

We asked 6 external participants to watch the movie and write down every time they thought there was a new event, given the following instructions: “Write down the times at which you feel like there is a major change in topic, location, time, etc. Each event should be more a less between 10 seconds and 3 minutes long. Don’t forget to write a small description of what was happening on that specific event.” With the participants’ boundary annotations we looked for those boundary time points that were consistent across observers. To find a statistical threshold of how many observers should coincide in a given time point to be different from chance in our data, we shuffled the number of observations 1000 times and created a null distribution of the resulting coincident time points. An α = 0.05 as a cutoff for significance indicated that boundary time points at which at least three observers coincided in (considering 3 seconds as possible window of coincidence) could not be explained by chance. The final human annotations model was composed of 38 events which is a number in accordance to the range found on the Chen et al. (2017) and Baldassano et al. (2017) studies.

### Verbal recall analysis

The audio files recorded during the free verbal recall were analyzed by a lab member which was a native Spanish speaker, using the list of events from the human annotations model. An event was counted as recalled if the participant described any part of the event and were counted as out of order if they were initially skipped and later described in the narrative.

### Finding event structure in the EEG data

To validate the event segmentation model extracted from human annotations on the EEG data collected from the primary sample in the study, we generated, for each of the individuals, a temporal correlation matrix computed by correlating the 29 electrodes with the same 29 electrodes for each of the time points. Next, we averaged the correlation values within each of the 38 events and ran a permutation test (N = 1000) with null boundaries picked by shuffling the temporal order of the events while maintaining their lengths. The within event correlation values were compared to the permuted values using an alpha of 0.05 as a cutoff for significance (see Figure 2b).

### Shared event neural patterns across individuals during movie viewing

Following previous fMRI findings (Chen et al., 2017), we examined whether neural patterns elicited by events during movie viewing were similar across individuals in our sample. To address this issue, we computed Movie-Movie correlations by comparing patterns of each event from one participant with the movie pattern for the same event averaged across the remaining participants. This resulted in an across-participants similarity analysis. To assess if the correlation values were statistically significant a permutation test (N = 1000) was computed, using an alpha of 0.05 as a cutoff for significance, by shuffling the event correspondence between the held-out and remaining participants.

### EEG pattern similarity within and across events

A similarity analysis was calculated for EEG neural activity before and after boundaries during movie viewing. The similarity analysis was performed at individual level, and included spatial (i.e., scalp voltages from all the 29 electrodes), and the temporal features, which were selected using a 200 ms sliding window of the resulting z-transformed signal obtained from averaging the EEG trials. The similarity analysis was calculated using Pearson correlation coefficients, which are insensitive to the absolute amplitude and variance of the EEG response.

The similarity analysis was computed on EEG segments of 10 seconds pre- and post-boundaries identified in the event segmentation model. To ensure that differences between before and after the boundary were not arising just due to intrinsic temporal contiguity properties of the EEG signal, we first split pre- and post-boundary 10-second EEG segments in two equal EEG vectors of 5 seconds. Thus, pre-boundary event correlations were performed between the interval -10 s to -5 s and the interval -5 s to 0 s before the boundary. Between-event correlations were performed between -5 s to 0 and 0 to 5 s, were 0 corresponded to the boundary. Post-boundary event correlations were performed on EEG data from the interval 0 to 5 s and the interval 5 s to 10 s after the boundary. Point-to-point correlation values were then averaged for each of the 3 conditions and differences were statistically compared by means of a repeated measures ANOVA including type of event as a 3-level factor (Pre-boundary, between-event, and post-boundary). Statistical significance was set at p < 0.05. A paired-sample t-test was used to test for statistical significance between condition pairs.

An EEG similarity analysis was also performed on 20-second windows of averaged EEG trial data as a function of whether events preceding a boundary were later recalled or forgotten. The 10 seconds EEG signal included 10 seconds before boundary and 10 seconds after boundary. Similarity analysis was implemented by correlating point-to-point the spatial EEG features surrounding the boundary. To account for differences between recalled and forgotten conditions a cluster-based permutation test was used (Maris and Oostenvelt, 2007), which identifies clusters of significant points in the resulting 2D matrix in a data-driven manner and addresses the multiple-comparison problem by employing a nonparametric statistical method based on cluster-level randomization testing to control the family-wise error rate. Statistics were computed for every time point, and the time points whose statistical values were larger than a threshold (p < 0.05, two-tail) were selected and clustered into connected sets on the basis of x,y adjacency in the 2D matrix. The observed cluster-level statistics were calculated by taking the sum of the statistical values within a cluster. Then, condition labels were permuted 1000 times to simulate the null hypothesis and the maximum cluster statistic was chosen to construct a distribution of the cluster-level statistics under the null hypothesis. The nonparametric statistical test was obtained by calculating the proportion of randomized test statistics that exceeded the observed cluster-level statistics.

### EEG evoked responses at boundary onset

Event-Related Potentials (ERPs) at boundary onset were calculated for each individual as a function of whether events were later recalled or forgotten in the free recall task. To obtain the ERPs, we first applied a low-pass filter of 12 Hz to the none-down-sampled EEG data. Then, epochs of -1000 to 2000 ms were chosen around each of the boundary time points determined from the event segmentation model. The pre-boundary interval (−100 to 0 ms) was used for baseline correction. ERP differences between recalled and forgotten conditions were investigated starting at 0 ms to 2000 ms after each boundary onset. Statistical significance of the differences between conditions was assessed by a cluster-based permutation test.

### Brain sources of ERPs

Low-Resolution Tomography Analysis (sLORETA) (Pascual-Marqui, 2002) was used to reconstruct the source space for ERP differences at boundary onset. This method performs localization inference based on images of standardized current density, which corresponds to the 3D distribution of electric neuronal activity that has maximum similarity (i.e. maximum synchronization), in terms of orientation and strength, between neighboring neuronal populations (represented by adjacent voxels). LORETA was calculated separately for each participant’s averaged ERP triggered by boundaries that followed events that were later recalled and forgotten. Source reconstruction for each condition was compared and results were displayed by means of t-values (paired t-test, one-tail).

### Reinstatement of EEG event patterns during free recall

We adapted the HMM approach used in Baldassano et al. (2017) and tested the extent to which it identified EEG-based latent-states during recall according to the event segmentation model constructed through human annotations during movie-watching.

The model is a variant of an HMM in which the latent variables are the event labels *s*_*t*_ for each timepoint *t* and the spatial signatures *m*_*k*_ (brain activity patterns across all EEG channels) for each event *k*. From the observed brain activities *b*_*t*_ we infer both *s*_*t*_ and *m*_*k*_. The model is set to assume that, for all the participants, the event starts in *s*_*1*_*=1* and ends with *s*_*T*_*=K*, where *T* is the total number of time points and *K* is the total number of events. We assume that in each time point we can either advance to the next state or remain in the same one, which results in a transition matrix where all elements are zero except for the diagonal and the adjacent off-diagonal.

An isotropic Gaussian model is used to compute the observation model so that the probability that a given observation, *b*_*t*_, is created by a state *s*_*t*_=*k* can be given by:

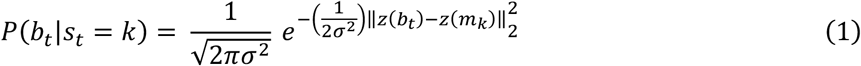

where z() denotes the z-score function. The z-scoring of the brain observations and the mean activity patterns result in a proportionality between the log probability of observing brain state *b*_*t*_ in an event with signature *m*_*k*_ and the Pearson correlation between *b*_*t*_ and *m*_*k*_ plus a constant offset:

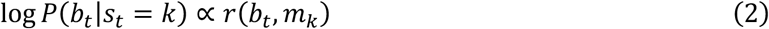

To ensure that all states are visited, the observation probabilities *P*(*b*_*t*_|*s*_*t*_ = *k*) are modified by setting *P*(*b*_F_|*s*_F_ = *k*) = 0, for all *k* ≠ *K* so that, on the final time point, only the final state *K* could have generated the data. To ensure that all possible event segmentations have the same prior probability, a dummy absorbing state *K+1* is created, so that the transition probabilities for state *K* are identical to those for previous states. We set *P*(*b*_*t*_|*s*_*t*_ = *K* + 1) = 0 so that this state cannot actually be used.

We used the mean EEG patterns of each of the events identified in the event segmentation model during movie watching to model the EEG data during recall. For each participant, the HMM was applied to the continuous recall EEG data to obtain a probabilistic assignment of latent event states consistent with the event segmentation model obtained during movie watching. The resulting probabilities *P*(*s*_*t*_ = *k*) were then used to identify the event transition points during recall, as timepoints when the most likely event changed. We then tested the extent to which HMM-based EEG patterns elicited during movie watching were similar to those estimated by HMM search during recall. As in the previous analysis, we ran an event-to-event correlation analysis between movie and recall and calculated an averaged correlation measure, as a proxy of the overall degree of similarity over the entire event segmentation model between the two sets of data. To assess for statistical significance at individual level, we created a null distribution by shuffling the movie events, then running the HMM on this shuffled order and finally computing the correlation between the shuffled movie events and the HMM-identified recall. This procedure was applied 1000 times to create a null distribution of findings and used to assess a statistical significance when compared with the real correlation values using an alpha of 0.05 as a cut-off.

## Author contributions

M.S., C.B. and L.F. designed the experiment. M.S. acquired the data. M.S. and L.F. analyzed the data. C.B. contributed to analytical methods. M.S., C.B. and L.F wrote the manuscript.

## Acknowledgements

This research study was supported by grants from the European Union – Erasmus+ program to M.S., the Spanish Government (PSI2016-80489-P) and the Catalan Government to L.F. (2017 SGR 1573).

## Supplementary Information

**Supplementary Table 1.** Distribution of recalled and forgotten events. We created a table of contingency for each of the participants to assess the possibility that recalled or forgotten events during movie-watching were non-uniformly preceded or followed by either recalled or forgotten events. Discarding this possibility is relevant to interpret our similarity (Figure 5a) and ERP (Figure 5b and c) findings at boundaries as specifically related to recall or forgotten in the later free recall task. For each participant, we performed a Fisher’s exact test to statistically assess for an unequal distribution of events. This analysis, however, resulted non-significant (p > 0.05) for all participants, thereby indicating that the distribution of recalled and forgotten trials was uniform.

|  | Events<br>Recalled | Events<br>Forgotten | Total |
| --- | --- | --- | --- |
| Events<br>Recalled | 11.57 (± 0.64) | 7.03 (± 0.20) | 18.60 |
| Events<br>Forgotten | 7.00 (± 0.21) | 11.40 (± 0.68) | 18.40 |
| Total | 18.57 | 18.43 | 37 |

**Supplementary Figure 1.**
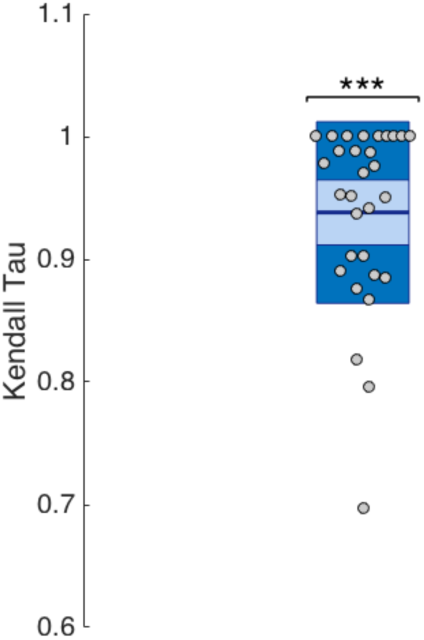
Preservation of event temporal order during free recall. To statistically assess whether the order of events during movie-watching was preserved during free recall, we computed Kendall rank correlation coefficients between each individual event temporal order and a simulated correct linear order. For all participants the Kendall tau coefficient was positive and close to 1, indicating that the encoded temporal order of the events was highly preserved during their recall. The central mark is the median, the edges of the box are the 25^th^ and 75^th^ percentiles. *** denotes p < 0.001.

**Supplementary Figure 2.**
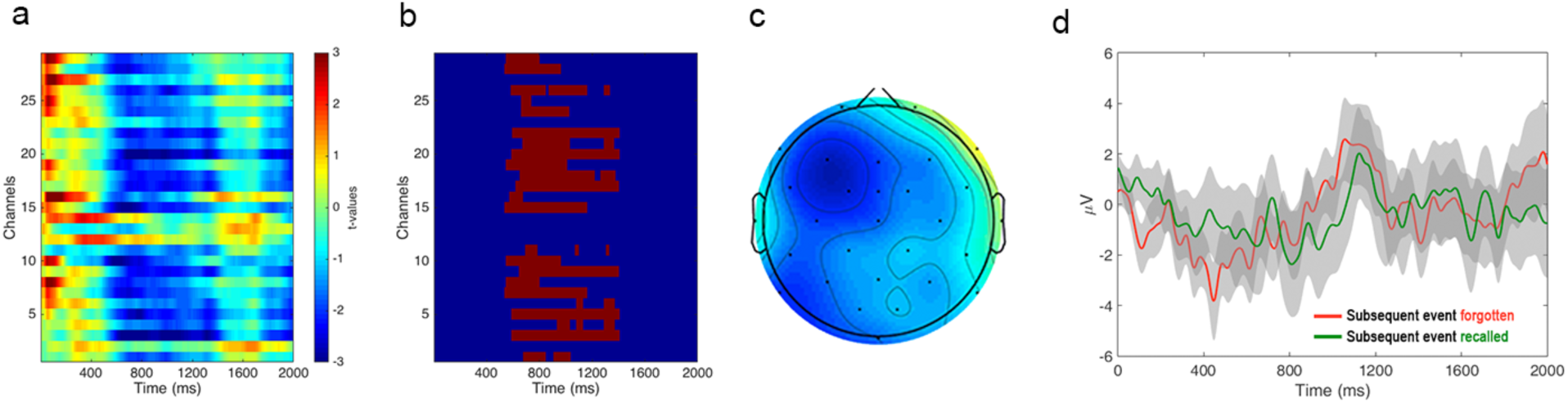
Event Related Potentials (ERPs) at boundaries. (**a**) Point-to-point ERP difference at boundary onset following events that were later recalled or forgotten. Differences are expressed in t-values (paired t-test). (**b**) Cluster of spatiotemporal contiguous points that resulted significant (p < 0.05) when the two ERP conditions were compared with a cluster-based permutation test (see Methods). (**c**) Scalp ERP representation of the ERP difference between the two conditions averaged over time-points within the significant cluster. (**d**) ERPs elicited at boundaries *preceding* events that were later recalled or forgotten. No differences were found between the two conditions (p > 0.05, cluster-based permutation test).

